# A rapid reversed-phase LC-MS method for polar metabolite profiling

**DOI:** 10.64898/2026.02.28.708719

**Authors:** Maxime De Neys, Jana K. Geuer, Sammy Pontrelli

## Abstract

Polar metabolites, including amino acids, nucleotides, phosphorylated metabolites, and central carbon intermediates, drive essential physiological processes but remain difficult to measure by high-throughput, reversed-phase LC-MS due to poor retention on conventional stationary phases. We developed a 3-minute reversed-phase LC-MS method and benchmarked T3-type C18 and pentafluorophenyl (PFP) chemistries for profiling 123 polar metabolites across acidic and mildly acidic conditions. The T3 phase operated under mildly acidic conditions provided the best overall performance, achieving the highest coverage, robust retention-time stability, and improved detection of phosphorylated metabolites. To strengthen compound annotation under ultra-short gradients, we combined the method with iterative data-dependent MS/MS, acquiring spectra for 86 of 123 metabolite mixture compounds without extending runtime. Retention times and peak shapes remained stable over 480 consecutive injections (mean CV of 1.7%) in *Escherichia coli* extract. Together, these results define a rapid, scalable workflow for profiling of polar and phosphorylated metabolites on standard instrumentation for high-throughput biological studies.

## Introduction

High-throughput LC-MS of highly polar metabolites remains a persistent analytical challenge, particularly for chemically and biologically important small molecules such as phosphorylated intermediates, organic acids, and nucleotides. Hydrophilic interaction chromatography (HILIC) is widely used to retain polar metabolites; however, long equilibration times, sensitivity to subtle changes in mobile phase composition and sample matrices, and retention time instability limit its suitability for high-throughput workflows^1–3^. At the other extreme, flow injection analysis (FIA) maximizes throughput by eliminating chromatographic separation entirely, but this approach exacerbates matrix effects, limits analyte discrimination, and reduces confidence in metabolite annotation^4^. Recent fast metabolomics methods using mixed-mode LC or HILIC coupled to ion mobility have demonstrated that short (≈3.5-5 min) gradients can still provide broad metabolome coverage, but typically at the cost of specialized hardware that is not universally available (i.e., quaternary or tertiary pumps or ion mobility-enabled mass spectrometers)^5,6^. Consequently, there remains a need for LC-MS methods that occupy a middle ground, offering short method times while retaining sufficient chromatographic separation to support robust metabolite profiling and annotation on standard UHPLC systems.

Conventional C18 stationary phases provide limited retention for highly polar compounds; however, recent advances in reversed-phase stationary phase design have expanded the applicability of reversed phase LC-MS to more polar analytes. Functionalized chemistries such as C18 T3-type^7^ and pentafluorophenyl (PFP) phases^8^ provide mixed-mode interactions, and stable retention in 100% aqueous mobile phases, enabling robust retention of polar compounds. Both column types have been shown to extend reversed-phase retention while preserving short equilibration times^3,7^. In parallel, the adoption of solid-core particles and reduced particle sizes has enabled higher separation efficiency and shorter gradients without prohibitive backpressure^9^. Despite these developments, the performance limits of such approaches under extreme throughput constraints, particularly for highly polar and phosphorylated metabolites, have not been systematically evaluated.

Here, we evaluate two complementary solid-core reversed-phase columns, a PFP phase and a T3-type C18 (T3) phase, across acidic and mildly acidic mobile phase conditions to assess their suitability for fast LC-MS analysis of highly polar metabolites. To develop a 3-minute sample-to-sample injection method, we benchmark retention time stability, peak shape, reproducibility, and analytical sensitivity for a panel of 123 polar metabolites spanning amino acids, organic acids, nucleotides, and phosphorylated intermediates.

MS/MS provides structurally informative fragmentation spectra that support metabolite identification. While the highest annotation confidence is achieved through direct comparison with analytical standards, spectral library matching substantially expands coverage^10^. Under ultra-fast chromatographic conditions, retention of highly polar metabolites is limit conventional data-dependent acquisition (DDA), which preferentially fragments the most intense precursor ions detected in MS1. However, short peak widths restrict the number of precursors that can be sampled, limiting MS/MS coverage and annotation depth. To overcome this limitation while preserving a 3-minute runtime, we implemented iterative DDA with dynamic exclusion^11^. By excluding previously fragmented precursors across repeated injections, precursor ions fragmented in earlier injections are excluded from subsequent runs. This enables progressive acquisition of MS/MS spectra from lower-abundance ions, increasing cumulative MS/MS coverage on representative samples across injections without extending per-sample analysis time during routine profiling^12^.

Central carbon metabolites are used as a chemically demanding test set to benchmark this method, as they encompass a broad range of highly polar analytes that challenge both chromatographic retention and ionization efficiency. Together, this work outlines practical design considerations for high-throughput reversed-phase LC-MS analysis of polar metabolites and provides a complementary alternative to HILIC- and FIA-based workflows for applications prioritizing analytical speed and scalability.

## Experimental Section

### Metabolite standards

All 123 standards were purchased from Sigma-Aldrich, weighed, and dissolved in a final concentration of 20 mmol/L per stock solution. From these stock solutions, a standard mix of 50 µmol/L per metabolite was created and stored in aliquots at -80°C.

### LC-MS conditions

LC-MS measurements were performed using an Agilent Infinity III stack either with an Agilent InfinityLab Poroshell 120 PFP column (2.1 mm x 50 mm, 1.9 μm) operated in reversed phase, with an InfinityLab Poroshell 120 PFP guard column (2.1 mm x 5 mm, 1.9 μm) at 30°C, or with a Waters CORTECS™ Premier T3 VanGuard™ FIT column (2.1 mm x 50 mm, 1.6 μm) operated in revered phase at 50°C. The autosampler was kept at 4°C and the injection volume was 5 μL. The injector needle was washed before each injection consecutively with three solutions for 3 s each to prevent carry-over: Methanol (MeOH, Fisher Scientific, USA)/H_2_O 50/50 (v/v), 2-Propanol (IPA, Biosolve, the Netherlands)/H_2_O (Milli-Q IQ 7000 purification system (MilliporeSigma, USA) 50/50 (v/v), and eluent A. Gradient elution was carried out at 1 mL/min with eluent A consisting of H_2_O/Acetonitrile (ACN, Fisher Scientific, USA) 100/0 (v/v) and eluent B consisting of ACN/H_2_O 90/10 (v/v), both containing either 0.1% formic acid (TCI Chemicals, Japan) (pH 3) or 0.1% ammonium acetate (Sigma-Aldrich, USA) buffer set to pH 5 using acetic acid (TCI Chemicals, Japan). After evaluating different higher organic compositions (70-80% B), the final gradient was as follows: 0 – 0.5 min 0% B, 0.5 – 0.9 min increased to 10% B, 0.9 – 1.5 min increased to 80% B, 1.5 – 1.98 min held at 80% B, 1.98 – 1.99 min returned to 0% B, and held for 1 min for re-equilibration. An Agilent Revident LC/Q-TOF was used for measurements, operated both in negative and in positive ionization modes with an m/z range of 20 – 1700 in 750 V stable mode. Source conditions were set to a gas temperature of 325 °C, drying gas of 10 L/min, nebulizer pressure of 30 psi, sheath gas temperature of 250 °C, and sheath gas flow of 11 L/min. The capillary voltage was set to 3500 V, with a nozzle voltage of 1000 V and octopole RF of 750 V. The capillary current was 0.24 µA and the chamber current (CA) was 1.7 µA. The fragmentor voltage was set to 110 V and the skimmer to 45 V.

### Iterative MS/MS

Iterative MS/MS experiments were performed on the pre-mixed standard mixture. The standard was thawed and diluted to a final concentration of 25 µmol/L. LC conditions were as described under LC-MS conditions exclusively on the T3 column. DDA was conducted on an Agilent Revident QTOF operated in both positive and negative electrospray ionization modes with source conditions as described under LC-MS conditions. A maximum of four precursors per cycle was selected using a narrow 1.3 m/z isolation window. An absolute intensity threshold of 5000 counts and a relative threshold of 0.03 % were applied for precursor selection. Charge state preference was restricted to singly charged ions and ions of unknown charge. Iterative MS/MS precursor exclusion was performed with a mass tolerance of ±20 ppm and a retention time exclusion window of ±0.5 min. Active exclusion was enabled after one MS/MS spectrum and released after 0.2 min. For MS only acquisition, data were acquired at 10 spectra/s with an acquisition time of 100 ms per spectrum. For MS/MS acquisition, a rate of 5 spectra/s was used with 200 ms per spectrum with the collision energy set to 20 eV.

### Data analysis

Data analyses were performed using Agilent MassHunter Qualitative/Quantitative Analysis software and R 4.5.1/R Studio 2025.09.2+418 with the packages tidyr^13^, dplyr^14^, and ggplot2^15^. MassHunter software was used to integrate peaks (Agile2 Integrator), fit polynomial calibration curves, and calculate retention times, full width at half maximum (FWHM), and signal-to-noise (S/N) ratios. Standards were measured for the following concentrations: 50, 25, 12.5, 10, 6.25, 3.125, 1.565, 0.781, 0.391, 0.195, 0.098, 0.049, and 0.024 µmol/L. Mean retention times (RT) were calculated using 20 replicate injections at 10 µmol/L. For the determination of the limit of detection (LOD) and limit of quantification (LOQ), 4 replicate injections were acquired at each concentration level, and levels with at least 3 successful replicate injections were retained. For each eligible level, the mean S/N ratio was calculated from the 3-4 replicates, and the LOD and LOQ were assigned as the lowest concentrations with mean S/N ≥ 3 and ≥ 10, respectively. Coefficient of variations (CV), LODs, LOQs, and retention factors (R_f_) were computed in R. The R_f_ was defined as the difference between standards’ mean retention times 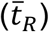 and the void time (t_0_), divided by the void time.

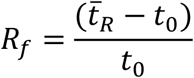

For iterative MS data files were processed using MSConvert (ProteoWizard)^16^, using the Peak Picking filter with vendor algorithm for MS1-2. Afterwards, the data and spectral libraries were imported into MZmine4^17^ and processed using a structured batch workflow. Mass detection was performed with noise levels of 500 (MS^1^) and 200 (MS/MS). MS^n^ tree building used an m/z tolerance of 20 ppm. Isotope detection (H,C,N,O,S) was applied with 20 ppm and maximum charge 1, selecting the most intense isotope. Features were aligned using the Join Aligner (15 ppm, 0.03 min RT tolerance, equal weighting). Correlation grouping used an RT tolerance of 0.03 min, minimum feature height 100 000, and an intensity threshold of 100. Ion identity networking was performed with 10 ppm intra-sample m/z tolerance, using an adduct library containing [M–H]^−^, [M+H]^+^, [M+Na]^+^, and [M+NH_4_]^+^. MS/MS spectral matching was performed using the publicly available libraries MassBank^18^, MassBank of North America (MoNA^19^), and GNPS^20^ in MZmine on merged representative MS/MS scans (per energy and merged across energies). Matching used 10 ppm tolerance for merging, 8 ppm for precursors, and 10 ppm for fragments. Precursor ions were removed prior to scoring. A minimum of 2 matched fragment ions and weighted cosine similarity with a minimum cosine score of 0.65 were required. After export, the annotations were matched against the list of standards in R4.5.2/R studio 2025.09.2+418 using the data.table package^21^.

### Bacterial growth conditions and extraction of metabolites

*Escherichia coli* BW25113 was stored at -80°C in glycerol stocks and streaked onto Luria-Bertani (LB) plates containing 2% (w/v) agar. A single colony was inoculated into 3 mL of liquid LB and grown overnight at 37°C with shaking at 200 rpm. The overnight culture was diluted 1:100 into fresh LB and incubated at 37°C, 200 rpm until mid-exponential phase (OD_600_ ≈ 0.5). Cells were harvested by centrifugation with subsequent removal of the supernatant. For intracellular metabolite extraction, the pellet was resuspended in cold extraction solvent consisting of ACN/MeOH/H_2_O, 40:40:20 (v/v/v) and incubated for one hour at -20°C. Intracellular metabolite extracts were separated by centrifugation, and the supernatant was collected and filtered through a 0.2 μm membrane prior to LC-MS analysis.

## Results & Discussion

To determine whether ultra-fast reversed-phase separations provide sufficient chromatographic structure for polar metabolites, we evaluated chromatographic performance under a 3-minute injection-to-injection workflow using two solid-core reversed-phase stationary, a PFP phase and a T3-type C18 phase, operated at pH 3 and pH 5 (**Figure 1**).

**Figure 1.**
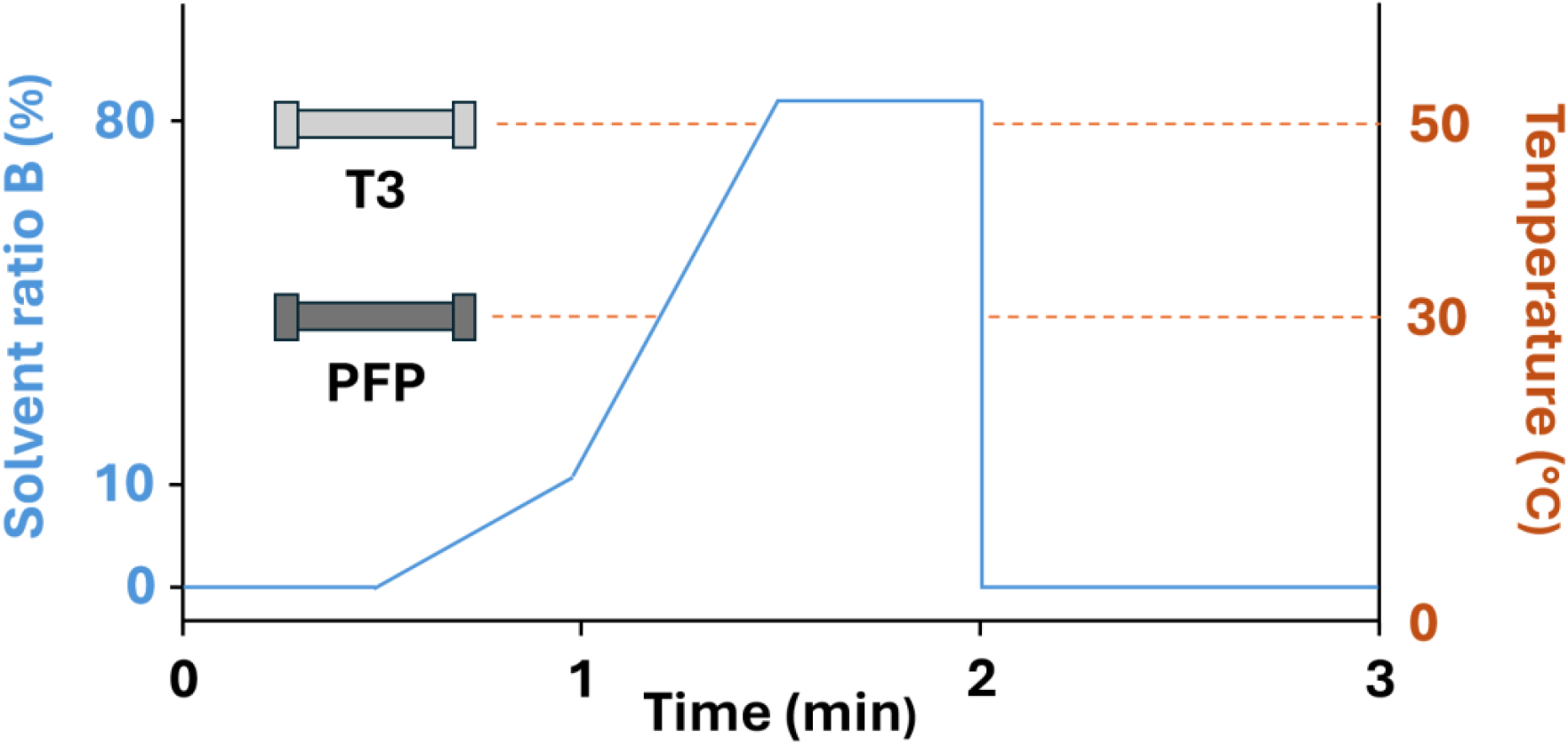
Reversed-phase gradient program. (Percentage of mobile phase B over time, blue) and column temperature configurations for the different columns, which were used for the 3-min LC-MS method (orange).

Both columns provided reproducible total ion chromatograms (TIC) (**Figure S1**) at a flow rate of 1 mL min^−1^ without exceeding instrument backpressure limits. The T3 column showed a greater increase in backpressure over time (0.05 bar/injection) relative to the PFP column (0.01 bar/injection), consistent with its smaller particle size (1.6 um compared to 1.9 um) and inlet frit (0.2 μm compared to 0.5 μm). Despite the short gradient, retention time precision was high with median RT CVs ranging from 1.05-1.48% across all column, pH, and ionization mode combinations **(Table S1)**. Chromatographic peaks remained narrow and well defined, with a median FWHM at 1.038 s. Together, these results demonstrate that even under extreme 3-minute gradient conditions, reversed-phase LC provides stable and reproducible analysis.

### Effect of column chemistry and pH on polar metabolite coverage

To assess how column chemistry and mobile phase pH influence analytical coverage, we quantified detected metabolites across all column, pH, and ionization mode combinations using a panel of 123 highly polar compounds. Across conditions, detected metabolites ranged from 69 to 114, indicating a strong dependence on both stationary phase and mobile phase composition (**Table S1, Figure 2A**). Negative ion mode consistently yielded higher coverage than positive mode. In negative mode, the T3 column outperformed the PFP at both pH values, with the highest overall coverage observed for T3 at pH 5 (114 metabolites), compared to the 82 metabolites for the best-performing PFP condition.

**Figure 2.**
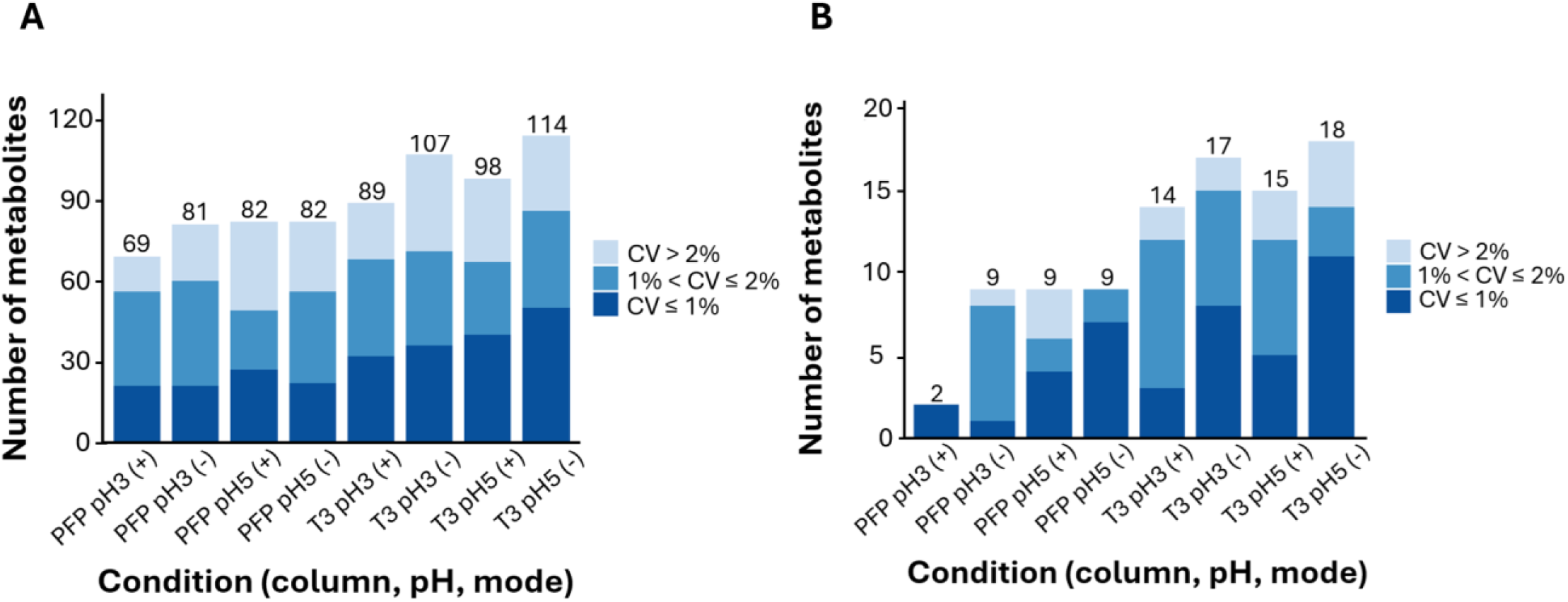
Effect of column chemistry, mobile phase pH, and ionization mode on metabolite coverage under 3-min gradient conditions. The number of detected metabolites across PFP and T3 columns operated at pH 3 and pH 5 in positive and negative ionization modes is shown for (A) the full polar metabolite panel (n = 123) and (B) the phosphorylated subset (n = 20).

Metabolites detected in negative mode were consistently observed as deprotonated species across all conditions. In positive mode, most compounds were detected as protonated species, although several highly oxygenated metabolites, predominantly carbohydrates and polyols, were preferentially or exclusively detected as sodium adducts (**Table S2**). Across all conditions, four metabolites (phenyl acetate, ethyl-3-methyl-2-oxobutyrate, guanosine 5’-triphosphate, and 2,3-diphospho-D-glucuronic acid) were not detected, likely reflecting poor ionization efficiency, instability or adsorption-related losses.

To contextualize coverage differences, analytical sensitivity was evaluated by estimating LOD and LOQ under each condition. Median LODs were comparable between T3 and PFP at pH 3 (0.0244 µmol/L in negative mode for both), but were lower for T3 at pH 5 (0.0977 µmol/L compared to 0.1953 µmol/L for PFP). LOQ trends followed a similar pattern, with T3 at pH 3 yielding the lowest values overall (0.0244 µmol/L) (**Table S3**).

Differences between stationary phases were most pronounced for phosphorylated compounds. Of the 20 phosphorylated metabolites included in the standard panel, 9 were detected in at least one PFP condition, whereas 18 were detected in at least on T3 condition (**Figure 2B**). This improved coverage is likely due to the bio-inert T3 hardware, which reduces analyte-metal interactions that disproportionately affect phosphate-containing metabolites^8,22^.

Mobile phase pH further modulated coverage, with pH 5 conditions generally enabling detection of a broader range of polar metabolites than pH 3 (**Figure 2**). This effect was most evident for phosphorylated compounds, which showed improved signal intensity and peak shape under mildly acidic conditions.

For example, adenosine 5’-triphosphate (ATP) exhibited a substantially sharper, more symmetric peak and increased signal intensity at pH 5 compared to pH 3 on the T3 column and was not retained on the PFP column (**Figure 3**). Similar trends were observed for other phosphorylated intermediates (**Table S4**). Retention time reproducibility followed similar trends. For the T3 column operated at pH 5 in negative mode, 50 metabolites exhibited RT CVs ≤1% and 86 exhibited RT CVs ≤2%, compared to 22 and 56, respectively, for the best-performing PFP condition. Although PFP resolved many compounds and has been shown to be effective for quantitative metabolomics in bacteria, providing good retention, sensitivity, and reproducibility^23^ our results, taken together, identify the Premier T3 column operated at pH 5 as the most effective condition for maximizing coverage of highly polar, particularly phosphorylated metabolites. As such, the T3 column operated at pH 5 was selected for all subsequent experiments.

**Figure 3.**
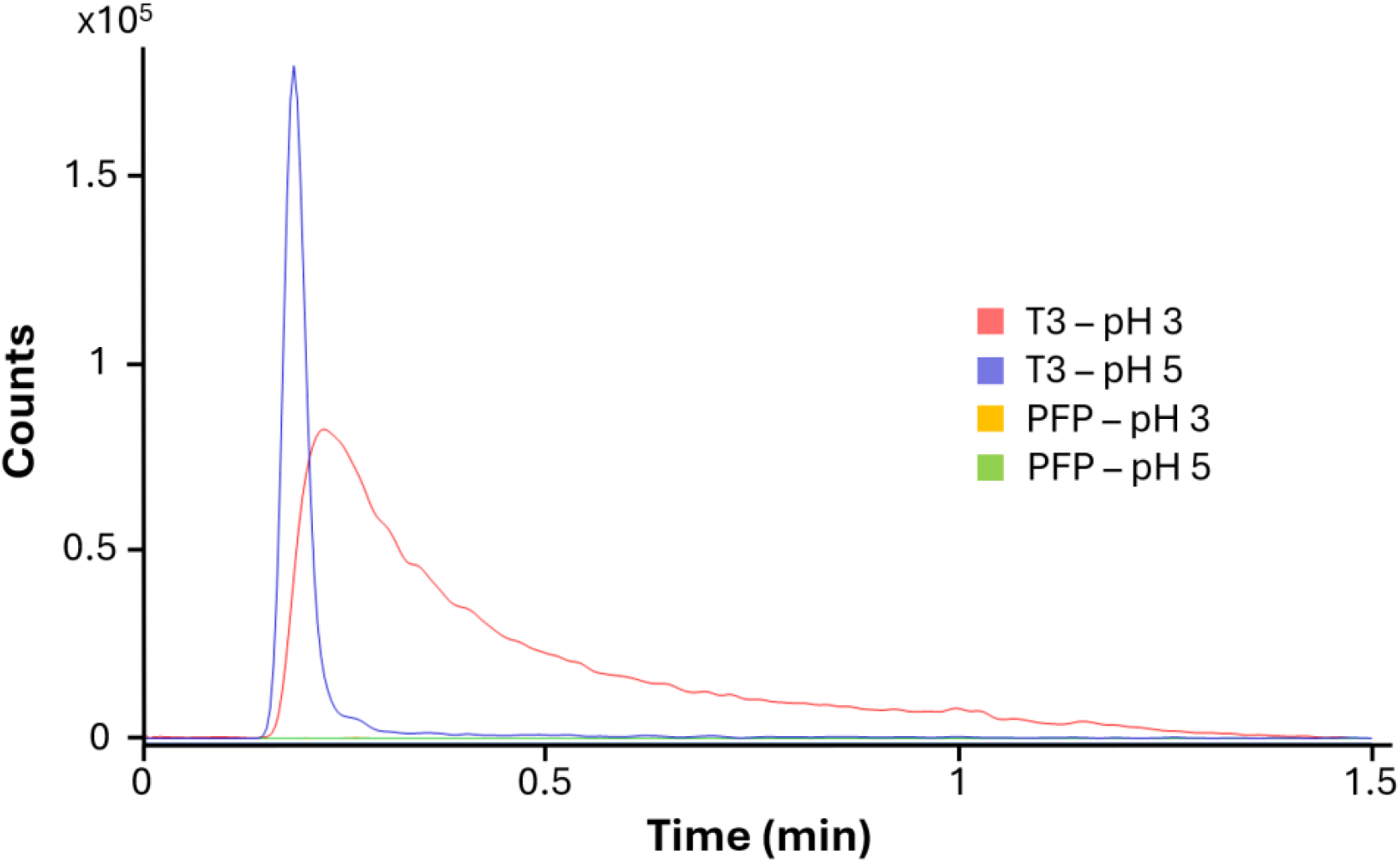
Effect of mobile phase pH on extracted ion chromatograms (EICs) of adenosine 5′-triphosphate. Smoothed EICs of adenosine 5’-triphosphate (50 µmol/L) acquired in negative ionization mode are shown for T3 at pH3 (pink) and pH5 (violet), and for PFP at pH3 (orange) and pH5 (green).

### Iterative MS/MS improves metabolite annotation despite limited retention

Despite the broad metabolite coverage achieved under this high-throughput workflow, the shortened gradient inherently limits chromatographic separation, constraining confident annotation of early, co-eluting compounds. To improve annotation confidence without compromising throughput, we employed repeated injections of a standard mixture to progressively acquire MS/MS spectra. Rather than relying on comprehensive MS/MS coverage within a single injection, this multi-injection strategy enables the progressive accumulation of fragmentation spectra for a larger fraction of features. In practice, this strategy allows MS/MS information to be accumulated from pooled samples, while routine profiling remains single-injection, with consistent retention times and peak shapes supporting metabolite annotation across large sample sets analyzed under identical conditions.

Using the T3 column, iterative MS/MS enabled annotation of 86 of the 123 metabolites through spectral library matching in MZmine 4. At pH 3, 72 metabolites were annotated. Of these, 24 were detected in both ionization modes, 21 were annotated exclusively in negative mode, and 27 were annotated exclusively in positive mode. At pH5, of 69 metabolites were annotated, of which 27 were picked up at both polarities, 22 in negative, and 20 in positive mode only (**Figure 4A**). Samples were iterated ten times. Across conditions, most metabolites were annotated within the first two iterations, where 35-65% of annotated metabolites captured in the first iteration. Additional MS/MS spectra were, however, acquired through the final iteration under pH3 positive mode conditions (**Figure 4B**). This differential dependence on pH is consistent with the ionization and detection trends observed above (**Table S1**), which influence precursor intensity and subsequent MS/MS acquisition.

**Figure 4.**
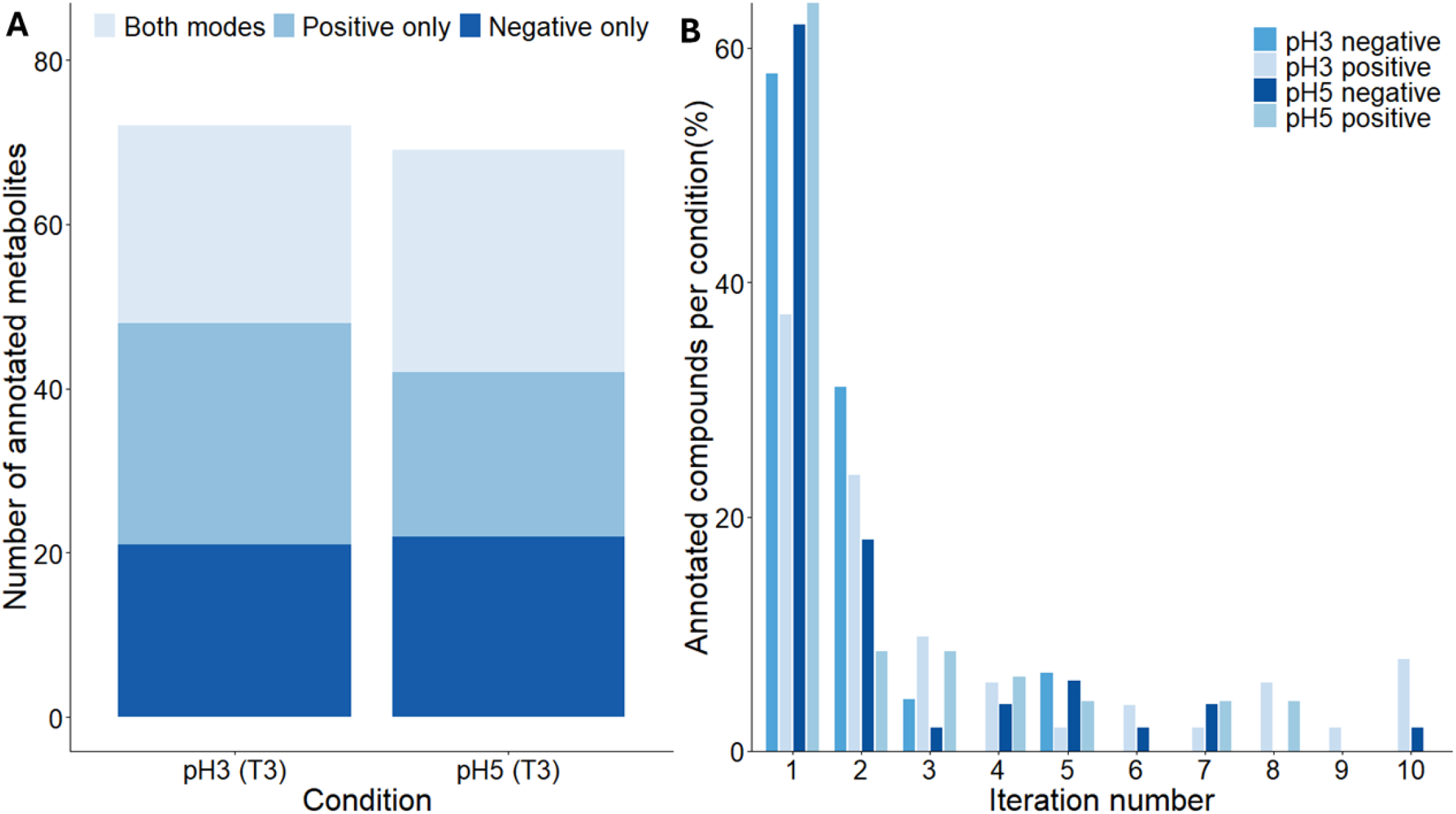
Annotations by iterative MS/MS. A: metabolites annotated based on spectral library matching by pH. In total, 69 metabolites were annotated at pH 5 and 72 at pH3 on a T3 column. Dark blue: compounds were annotated in both positive and negative mode. blue: compounds were only annotated in positive mode, bright blue: compounds were only annotated in negative mode. B: annotated compounds by number of iteration. Dark blue: Annotations at pH5, negative mode, medium blue: annotations at pH3, negative mode, blue: annotations at pH5 positive mode, light blue: annotations at pH3, positive mode.

### Method robustness and annotation performance in a complex biological matrix

To evaluate method robustness and annotation performance in a realistic sample matrix, we analyzed an *Escherichia coli* intracellular extract using the optimized T3 column operated at pH 5 in negative mode. The extract was injected repeatedly over 480 consecutive injections under identical chromatographic conditions to assess retention time stability and peak shape reproducibility over extended use.

Across the injection series, 94 out of the 123 monitored metabolites were detected in more than 95% of injections, indicating robust detection in complex *E. coli* matrix (**Table S5**). Median RT CVs across detected features were 1.7%, with 21 metabolites exhibiting RT CVs ≤1% and 51 metabolites exhibiting RT CVs ≤2% over the full injection sequence. Retention times in matrix closely matched those of neat standards, with a median ΔRT of 0.168 s across 94 metabolites. Peak shapes remained consistent throughout the run, with a median FWHM of 0.95 s across detected metabolites and no systematic broadening or tailing observed, indicating stable chromatographic performance under prolonged high-throughput operation. Backpressure increased gradually over the injection series but remained within acceptable operating limits, consistent with expectations for extended use of short, small-particle columns. Together, these results demonstrate stable chromatographic performance and metabolite detectability in complex biological matrices over extended high-throughput operation.

## Conclusions

By benchmarking two reversed-phase chemistries across mobile phase pH conditions, we show that the T3 column operated at pH 5 provides the most effective balance of coverage, RT reproducibility, and peak shape for polar and phosphorylated metabolites under high-throughput constraints, which aligns with other findings, which identified T3 as the best performing column in terms of chromatographic retentivity, peak shape, MS sensitivity and resolution in a comparative study on polar stress biomarkers^24^. The bio-inert T3 hardware minimizes secondary metal interactions, which likely further contributes to performance for phosphate-containing metabolites, which is a common issue in systems with many stainless steel surfaces^22^. We further demonstrate that repeated injections can be used to progressively acquire MS/MS spectra, substantially increasing annotation coverage without compromising throughput. While this strategy improves annotation confidence under short-gradient conditions, co-elution of matrix components is unavoidable, and ion suppression effects cannot be fully eliminated. Accordingly, similar to FIA-based workflows, the method described here is primarily intended for high-throughput metabolite profiling rather than absolute quantification.

Alternative strategies for rapid metabolome coverage have combined zwitterionic HILIC with ion-mobility separation^25^ or employed mixed-mode stationary phases with ternary gradients^26^ to achieve 3.5-5 min separations and broad metabolome coverage. While effective, these approaches require specialized hardware, including ion-mobility mass spectrometers or non-standard LC pump configurations (i.e., ternary or quaternary systems), which can limit routine adoption. By comparison, the approach presented here profiles highly polar and phosphorylated metabolites using a T3 reversed-phase column operated with a simple binary gradient on conventional LC-MS instrumentation, supporting accessibility and ease of deployment across laboratories.

Together, this work addresses practical design considerations for fast reversed-phase LC-MS analysis of polar metabolites and establishes a framework for balancing throughput, robustness, and annotation confidence. Importantly, the method described here deliberately focuses on the early portion of the reversed-phase gradient, where highly polar metabolites elute. In real biological samples, less polar and nonpolar metabolites would be expected to elute later in the gradient and were not evaluated in the present study. As such, this method provides a flexible and analytically grounded option for studies that require rapid, scalable metabolite profiling.

## Author information

### Author Contributions

The manuscript was written through contributions of all authors. All authors have given approval to the final version of the manuscript. ‡These authors contributed equally.

## Acknowledgements

We thank Els Meert for preparation of metabolite standards. This work was supported by VIB-KU Leuven Center for Microbiology and FWO Odysseus Type II funding (Grant number G0AT025N).

