## Supplemental information for "A rapid reversed-phase LC-MS method for polar metabolite profiling"

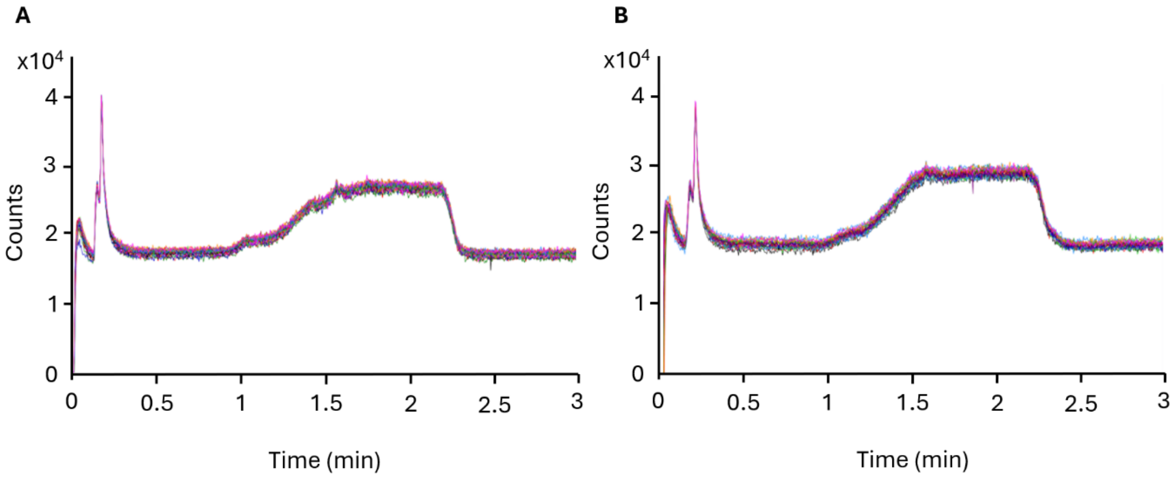

**Figure S1. Total ion chromatogram (TIC) reproducibility under reversed-phase conditions at pH 5 in negative ionization mode.** Overlayed TICs from 20 replicate injections (10 µmol/L) are shown for (A) the T3 column using a 20-metabolite standard mixture and (B) the PFP column using a 103-metabolite standard mixture.

**Table S2. Summary of detected ion species in positive mode across column chemistries and pH conditions.** Metabolites detected exclusively as sodium adducts are listed for each condition.

| Conditions | Total detected metabolites | Detected as protonated ions | Detected exclusively as sodium adducts |
|---|---|---|---|
| PFP pH3 | 69 | 57 | **12** (2′-Deoxyuridine, D-(−)-fructose, D-(−)-ribose, D-(+)-galactose, D-(+)-glucosamine hydrochloride, D-(+)-glucose, D-(+)-maltose monohydrate, D-(+)-mannose, D-(+)-trehalose dihydrate, D-(+)-xylose, myo-inositol, sucrose) |
| PFP pH5 | 82 | 71 | **11** ((±)-3-Methyl-2-oxovaleric acid sodium salt, 2′-deoxyuridine, D-(−)-fructose, D-(−)-ribose, D-(+)-galacturonic acid monohydrate, D-(+)-glucose, D-(+)-trehalose dihydrate, D-(+)-xylose, D-glucuronic acid sodium salt monohydrate, myo-inositol, shikimic acid) |
| T3 pH3 | 89 | 77 | **12** (2′-deoxyuridine, D-(−)-fructose, D-(−)-ribose, D-(+)-galactose, D-(+)-glucosamine hydrochloride, D-(+)-glucose, D-(+)-maltose monohydrate, D-(+)-mannose, D-(+)-trehalose dihydrate, D-(+)-xylose, myo-inositol, sucrose) |
| T3 pH5 | 98 | 83 | **15** ((±)-3-methyl-2-oxovaleric acid sodium salt, 2′-deoxyuridine, 3-hydroxy-1-propanesulfonic acid sodium salt, 3-hydroxyglutaric acid, D-(−)-fructose, D-(−)-ribose, D-(+)-galacturonic acid monohydrate, D-(+)-glucose, D-(+)-trehalose dihydrate, D-(+)-xylose, glycerol, glyoxylic acid monohydrate, L-valine, myo-inositol, shikimic acid) |

**Table S3. Median LOD and LOQ for detected metabolites across conditions** (column, pH, ionization mode)

| **Conditions** | **LOD in positive mode (µmol/L)** | **LOD in negative mode (µmol/L)** | **LOQ in positive mode (µmol/L)** | **LOQ in negative mode (µmol/L)** |
|---|---|---|---|---|
| PFP pH3 | 0.0244 | 0.0244 | 0.0488 | 0.0488 |
| PFP pH5 | 0.0488 | 0.1953 | 0.0488 | 0.1953 |
| T3 pH3 | 0.0244 | 0.0244 | 0.0244 | 0.0244 |
| T3 pH5 | 0.0488 | 0.0977 | 0.0977 | 0.0977 |

**Table S4. Mean full width at half maximum (FWHM) values for 20 phosphorylated metabolites under T3 conditions at pH 3 and pH 5 in positive and negative ionization modes.** Mean FWHM values were calculated from 20 replicate injections of a 10 μmol/L standard mixture under identical chromatographic conditions.

| Compounds | Mean FWHM* (second) | | | |
|---|---|---|---|---|
| | pH3 positive mode | pH3 negative mode | pH5 positive mode | pH5 negative mode |
| 2,3-Diphospho-D-glyceric acid | / | / | / | / |
| 6-Phosphogluconic acid | 0.908 | 1.113 | 0.621 | 0.573 |
| Adenosine 3′,5′-cyclic monophosphate | 0.855 | 0.896 | 1.022 | 0.894 |
| Adenosine 5′-diphosphate | 3.201 | 3.215 | 1.093 | 0.849 |
| Adenosine 5′-monophosphate | 0.911 | 1.070 | 0.959 | 0.843 |
| Adenosine 5′-triphosphate | / | / | 9.925 | 7.282 |
| D-Fructose 1,6-bisphosphate | 1.417 | 3.161 | / | 0.705 |
| D-Fructose 6-phosphate | / | 1.067 | / | 1.111 |
| D-Ribose 5-phosphate | 0.466 | 1.044 | 1.247 | 1.585 |
| Dihydroxyacetone phosphate | 0.622 | 1.161 | 1.253 | 1.696 |
| Glucose-6-phosphate | 1.216 | 1.067 | 1.090 | 1.111 |
| Glycerol phosphate | 0.776 | 0.99 | 0.526 | 2.183 |
| Guanosine 3′,5′-cyclic monophosphate | 1.825 | 2.663 | 1.133 | 0.915 |
| Guanosine 5′-diphosphate | / | 4.074 | 1.568 | 1.644 |
| Guanosine 5′-monophosphate | 0.884 | 0.950 | 0.841 | 0.793 |
| Guanosine 5′-triphosphate | / | / | / | / |

| | | | | |
|---|---|---|---|---|
| β-Nicotinamide adenine dinucleotide 2′-phosphate | 2.782 | 3.767 | 0.920 | 0.866 |
| Phospho(enol)pyruvic acid | 0.672 | 1.597 | 0.608 | 0.486 |
| α-D-Glucose 1-phosphate | 1.216 | 1.068 | 1.090 | 1.111 |
| β-Nicotinamide adenine dinucleotide 2′-phosphate, reduced | / | 2.340 | / | 1.059 |
